# Transfer Entropy Based Causality from Head Motion to Eye Movement for Visual Scanning in Virtual Driving

**DOI:** 10.1101/2022.10.10.511531

**Authors:** Runlin Zhang, Qing Xu, Zhe Peng, Simon Parkinson, Klaus Schoeffmann

## Abstract

Visual scanning is achieved by eye movement control for visual information acquisition and cognitive processing, which plays a critical role in undertaking common sensorimotor tasks such as driving. The specific coordination of the head and eyes, with head motions temporally preceding eye movements, is an important human behavior to make a key contribution to goal-directed visual scanning and sensorimotor driving. In this paper, we put forward a proposal of philosophy that this specific coordination of the head and eyes essentially indicates a unidirectional causality from head motion to eye movement. We propose to investigate transfer entropy for defining a quantitative measure of this unidirectional head-eye causality. A normalized version of the proposed causality measure is introduced for taking a role as an assessment proxy of driving. The plain transfer entropy-based definition has shown its statistical significance as the measure of causality and, the normalized version has demonstrated its good effectiveness for the evaluation of driving performance, with the verification in virtual reality-based psychophysical studies. This paper successfully suggests that the quantitative exploitation of causality based on the specific coordination of the head and eyes offers an effective approach to behaviometrics of visual scanning and sensorimotor activity.

**Author summary:** The coordination of head and eyes always exists in everyday sensorimotor driving tasks. Specifically, in goal-directed tasks, preparatory head motions guide eye movements to obtain and process relevant visual information for interacting with the surrounding environment. That is, the specific coordination of head and eyes involving head motions temporally preceding eye movement provides a mechanism for drivers to rely on prior knowledge for performing the tasks. As a matter of fact, this specific coordination of head and eyes essentially indicates, theoretically, a unidirectional causality from head motion to eye movement, leading to our proposal of causality philosophy. In this paper, an information-theoretic tool, transfer entropy, is exploited to capture the complex relationship between head motion and eye movement for obtaining the proposed measure of unidirectional causality. Furthermore, considering that the specific coordination of the head and eyes reflects the attention and cognitive state affecting the performance of sensorimotor tasks, we develop a normalized unidirectional causality measure as a proxy for the evaluation of driving performance. Psychophysical studies for goal-directed driving tasks are conducted based on virtual reality experimentation. Extensive results demonstrate a statistically significant correlation between the proposed normalized measure of causality and driving performance, which may provide a new and effective avenue for behaviometric applications. Practically, the merit of our proposed causality philosophy is that it is simple but effective, for obtaining an evaluation of the attentional and cognitive processes in driving tasks.

## Introduction

Visual scanning performed by rotating the eyes (the so-called eye movement) is important for human environment interactions [1, 2]. The investigation of visual scanning provides a fundamental window into the nature of visual-cognitive processing while performing naturalistic sensorimotor tasks such as walking and driving [3]. An activity of eye movement and visual scanning, which is composed of a sequence of gaze shifts, can be typically understood based on the technique of eye tracking [4–6]. According to the eye movement methodology [4–6], gaze point is an information element captured by eye tracker and, gaze points in some range and duration constitute a fixation; the gaze shift from one fixation to another represents a saccade. The understanding of eye movement and visual scanning provides an effective probe of human activity and performance in a sensorimotor task [1, 2, 7–9].

As has been largely accepted, a sensorimotor task, as a whole, is executed as a “top-down” goal-directed activity for meeting the task requirements [1, 2]. Gaze and eye movement are strongly directed based on the task goals, rather than on the “bottom-up” stimuli or irrelevant distractors [1, 2]. Therefore, firstly, the goal-directed driving is taken as the topic of our paper. Secondly, the underpinning mechanism of visual scanning and visual-cognitive processing essentially includes the coordination of head and eyes in the procedure of performing the sensorimotor tasks [10–14]. Indeed, eye movements larger than 15°, which are easily and usually happened, are successfully achieved based only on the synergistic coordination of head and eyes [7, 14]. Furthermore, both the head motion (namely, rotations of the head) and eye movement are usually and always involved in the visual scanning and visual-cognitive processing during performing a goal-directed sensorimotor task [1, 2, 7–9]. For example, the specific coordination of head and eyes, namely the preparatory head motion earlier than the eye movement, is involved to carry out soundly the curve driving and/or lane changing [12, 15]. As another example, even though a person performs a sensorimotor task that is to smoothly keep proceeding straight on a road consisting of straight sections, the performer uses this specific coordination to keep along an “imaginary” straight line [8, 16]. In this case, due to some possible factors involving the human, vehicle and environment, it is very hard for the performer to maintain the trajectory of vehicle strictly straight and thus, gaze shifting and fine tuning of the vehicle heading are always introduced for fixing on a visual target and also for keeping the vehicle on an imaginary straight line [17, 18]. Notably, what is stated above exactly means that the specific coordination involving head motions temporally preceding eye movements, called here the *specific head-eye coordination*, exists as a fundamental human behavior and mainly contributes to the visual scanning and visual-cognitive processing during a goal-directed sensorimotor task [11–14]. In the meanwhile, it should be pointed out that the coordination of head and eyes, with eye movements temporally preceding head motions, does play little role in the goal-directed sensorimotor tasks [11–14]. Further, the usual and always involvement of this *specific head-eye coordination* emerges strongly, no matter whether irrelevant distracting stimuli exist and no matter how many distractors present in the environments [1, 2, 7–9]. In other words, if there exists no irrelevant distractor, the *specific head-eye coordination* happens naturally [7, 9, 19]. If irrelevant distractors appear, a sensorimotor task is performed in a goal-directed fashion and distractors are ignored on purpose [20], then the *specific head-eye coordination* still contributes mainly to the task performing process [1, 2, 7–9]. Thirdly, the *specific head-eye coordination* itself does reveal information about the attention and cognitive state affecting the performance of sensorimotor tasks [12–14, 21, 22]. As a result, the quantitative characterization of the *specific head-eye coordination* and its quantitative influence on the driving performance will be centrally considered for the sake of this paper.

### The Philosophy of a Unidirectional Head-eye Causality

Eye movement and head motion data are observed as time series of gaze points ⟨*X*_t_⟩ and of head poses ⟨ *Y*_t_⟩, respectively, labelled by a sequential time index *t* = …, 1,2, … In this paper, stochastic processes, as usually used as natural representations for complex and real-world data [23], are introduced to model the time series data of eye movement and head motion, denoted by variables *X* and *Y*, respectively.

In the procedure of performing sensorimotor tasks, the observation of eye movement *X*_t_ can be intuitively considered as, in terms of some probability, being predicted by its past *X*_t−1_, for conducting the visual scanning activity [24]. In fact, the prediction of the eye movement time series can be added by head motion data, as the coordination of head and eyes always happens [11–14]. Notice that the systematic and comprehensive interplay happens between different body components, such as brain, visceral organs, head and eyes, in the visual-cognitive processing for the fulfillment of sensorimotor tasks [25, 26]. As a prominent example, *the specific head-eye coordination* mentioned above renders an important kind of interplay [11–14]. Notably, this specific coordination actually implies that, the past of head motion *Y*_t−1_ helps predict the current observation of eye movement *X*_t_. That is to say, formally, the probabilistic predictivity of *X*_t_ is added by *Y*_t−1_.

According to the essential concept behind the causality definitions by Wiener [27] and Granger [28], the Wiener&Granger causality (causality, for concise presentation, unless otherwise specified in this paper) from one stochastic process *Y* to another *X*, measures the probabilistic predictivity of *X*_t_ increased by *Y*_t−1_ [27]. And obviously, it is reasonably certain that *the specific head-eye coordination* essentially involves a kind of causality. Despite strong evidence that *the specific head-eye coordination* universally manifests as a typical trait of the goal-directed and task-driven behavior in visual scanning and sensorimotor driving [12, 14], it is unclear whether this *specific head-eye coordination* indicates a formal concept of causality. To investigate further, in this paper we propose a philosophy that the *specific head-eye coordination* under consideration can be characterized by a unidirectional head-eye causality, from head motion to eye movement. In fact, the proposed unidirectional head-eye causality not only provides an explainable determinant for the *specific head-eye coordination*, it also gives a kind of quantitative definition of this specific coordination.

Notice that, although the research on the dynamics of the coordination of head and eyes in the visual scanning and visual-cognitive processing has attracted a lot of studies recently [10–12, 14], there is no quantitative measure on this coordination. And in particular, quantitative, general and normative definition of causality based on the *specific head-eye coordination*, with head motions temporally preceding eye movements, for the sake of deep theoretical understandings, has never been touched. Transfer entropy [29], as an information theoretical tool in the sense of *Shannon* [30], is proposed here to be utilized for defining the measure of unidirectional head-eye causality. Due to the good properties of model-free and non-parametric manners involved in transfer entropy [23], the complexity and non-linearity of the time series data of eye movement and head motion [12] can be handled well.

It should be emphasized that the *specific head-eye coordination* offers a mechanism for visual scanning and for visual-cognitive processing [11], and furthermore, this specific coordination indicates an attentional state affecting the performance of goal-directed sensorimotor tasks [12–14, 21, 22]. Thus it is clear that the *specific head-eye coordination* definitely does have a direct correlation with the driving performance. However, little research has examined whether the quantitative measurement of this specific coordination itself behaves as an evaluation (proxy) of driving performance. In this paper we propose that a normalized version of the proposed unidirectional causality from head motion to eye movement, which gives a representation of the quantitative strength of the *specific head-eye coordination*, should furnish a proxy for the task performance.

To address the aforementioned gaps, we have accomplished our proposals and, have succeeded in obtaining two new quantitative causality measures for characterizing the *specific head-eye coordination* and driving performance, respectively. It is worthy to point out that the proposed transfer entropy based causality methodology provides a new point of entry for the investigation and discovery of a quantitative measurement for human behavior. The main contributions of this paper are listed as follows:

1. A head-eye causality philosophy is proposed to obtain a Unidirectional Causality Measure (*UCM*) for quantitatively evaluating a specific coordination of head and eyes during visual scanning, this *specific head-eye coordination* involves head motions temporally preceding eye movements. It is noted that, to the best of our knowledge, we originally define, formally and abstractly, the *specific head-eye coordination* in sensorimotor driving as a unidirectional causality from head motion to eye movement. Theoretically, this quantitative, general and normative definition of unidirectional head-eye causality can be considered as an essential and significant advance.

2. A Normalized Unidirectional Causality Measure (*NUCM*) is proposed for the purpose of being quantitatively compatible with the performance of sensorimotor driving. *NUCM* measure works well as a proxy of driving performance, verified by the psychophysical studies conducted in our paper. And this is a great benefit for application of behaviometrics.

3. As far as we know, the two proposed measures (*UCM* and *NUCM*) are the first discovery to successfully exploit transfer entropy for obtaining the unidirectional head-eye causality, as a determinant of the *specific head-eye coordination*, in the context of performance evaluation of sensorimotor tasks.

### Paper Organization

This paper is organized as follows. Firstly, related works are presented in Section *Related Works*. We then describe the proposed methodology for the new causality measures in Section *The Proposed Methodology for New Causality Measures*. The experiment conducted is detailed in Section *Experiment*, followed by the results and discussions in Section *Results and Discussions*. Finally, we present the conclusion and future works in Section *Conclusion and Future Works*.

## Related Works

### Wiener&Granger Causality and Transfer Entropy

The concept behind classic Wiener&Granger causality is one of the fundamental frameworks for defining causality [27, 28]. The abstract idea of Wiener&Granger causality is that, for two random variables *P* and *Q*, if the past of *Q* adds the predictivity of current *P*, then it is said that there is a causality from *Q* to *P*. The prediction idea by Wiener&Granger has been popularly accepted as a valuable and workable approach to defining causality [23], despite the causality definition based on prediction being under debate when compared with that on intervention proposed by Pearl [31].

Transfer entropy, basically as a measure of complexity, is a well-known way for quantifying the directional information flow between time series [29]. Transfer entropy is considered as a non-parametric and model-free version of Wiener&Granger causality, being capable of handling complex and non-linear time series [23]. Given random variables *P* and *Q*, transfer entropy from source *Q* to target *P* is defined as follows [23]:

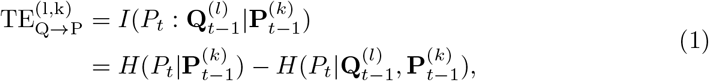

where *P*_t_ and *Q*_t_ are the observations of variables *P* and *Q* at time *t* respectively, 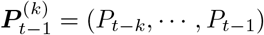 and 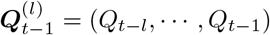 are the temporally ordered histories of target and source variables respectively, and *H*(·|·) and *I*(· : ·) represent respectively conditional entropy and mutual information. Here *l* and *k* are the so-called history lengths of 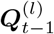and of 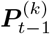 respectively. Notice that the information flow from *Q* to *P* obtained by 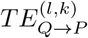 tries to take out the influences of the past of *P*.

Obviously, transfer entropy is asymmetric. Because 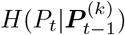 is no smaller than 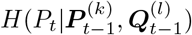, transfer entropy is nonnegative. Considering that 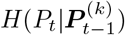 and 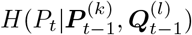 are nonnegative, 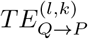 takes 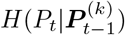 as the maximum.

### Coordination of Head and Eyes

Recently, many research studies have suggested the large popularity of the coordination of head and eyes in human activities [13, 14, 32, 33], for example in driving [14, 32] and motor control [13, 33].

The coordination of head and eyes always exists in our behavioral activities, particularly when a relatively large attentional shift is about to occur [1, 2, 7–9]; as a matter of fact, this coordination emerges as long as the eye movement is bigger than 15° [7, 14]. Specifically, the coordination of head and eyes is necessary because eye movements could selectively allocate the available attentional resources to task relevant information, and head motions could accommodate the limited field of view of the eyes [8, 9]. That is, head motions and eye movements are synergistic, especially temporally, for visual scanning and visual-cognitive processing [11]. Basically, head motions are followed by eye movements (namely, the preparatory head motion earlier than the eye movement) during sensorimotor tasks, because the observer usually has prior and “top-down” knowledge attaining attentional shift for goal-directed modulation [11–14, 34].

Note that the *specific head-eye coordination* involving head motions temporally preceding eye movements (rather than the coordination with eye movements temporally preceding head motions), has been definitively accepted as a main coordination of head and eyes [11–14], in the sense of statistics, in the goal-directed human activities. In fact, no matter whether irrelevant distracting stimuli exist and no matter how many distractors appear, the *specific head-eye coordination* principally contributes to goal-directed modulation during sensorimotor tasks [1, 2, 7–9]. When irrelevant visual distractors appear, performers allocate attention based on their goals and inhibit the effects of distractors [20], resulting in the preparatory head motion earlier than the eye movement. Even when no distraction is present, the *specific head-eye coordination* behaves naturally for humans [7, 9, 19]. In addition, the point here is that the directional coordination of head and eye movements (namely, the *specific head-eye coordination*) itself, does possess information about the performer’s attentional and cognitive state, affecting the task performance [12–14, 21, 22].

### Complexity Measures for Visual Scanning

In this paper, the complexity measures based on information entropy, which have been used for the assessment of visual scanning efficiency, are introduced.

The entropy rate can be identified by multiplying the summation of inverse transition durations and the normalized entropy of fixation sequence together [35]. Entropy of fixation sequence (*EoFS*) is *Shannon* entropy of the probability distribution of fixation sequences [36]. Gaze transition entropy (*GTE*) [37] is defined as a conditional entropy based on the probability transition between *Markov* states (namely, the areas of interest (*AOI* s)). Stationary gaze entropy (*SGE*) [37] gives the *Shannon* entropy based on an equilibrium distribution of *Markov* states. A latest technique called time-based gaze transition entropy (*TGTE*) [14], which uses time bins to realize the idea of *GTE*, is proposed for handling visual stimuli with dynamic changes.

## The Proposed Methodology for New Causality Measures

### A Unidirectional Causality Measure (*UCM*)

As discussed in Section *The Philosophy of a Unidirectional Head-eye Causality*, the *specific head-eye coordination* can be exploited as a measure of the unidirectional causality from head motion to eye movement (namely, *UCM*) and, transfer entropy is selected to investigate this causality due to its suitability. Following (1), transfer entropy from head motion *Y* to eye movement *X, TE*_Y →X_, is defined as:

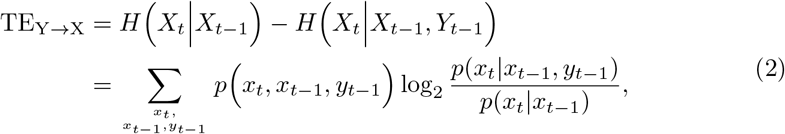

note in this paper, the history lengths of *X* and *Y* are both taken as 1, as usually done in literature [23]. Other possible options of the history length are outside of the paper scope but will be considered in the near future. Here *p*(·) and *p*(·|·) denote the (conditional) probability distributions of gaze (*x*_t_) and head (*y*_t_) data. And similarly, *TE*_X→Y_, transfer entropy from eye movement to head motion, is given as follows:

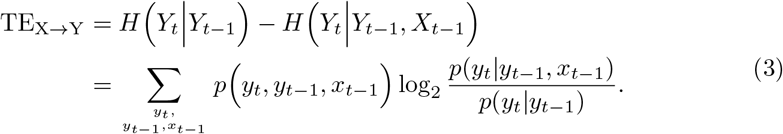

Notice that the more predictivity of current *X* added by the past of *Y*, the larger *TE*_Y →X_ is, the larger the causality from *Y* to *X*. Analogously, the more predictivity of the current *Y* added by the past of *X*, the larger *TE*_X→Y_ is, the larger the causality from *X* to *Y*. In this case, the unidirectional causality from head motion *Y* to eye movement *X* can be, effectively and firmly, detected, discovered and defined as *TE*_Y →X_ minus *TE*_X→Y_,

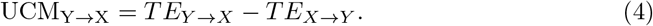

The rationale of *UCM*_Y →X_, as a unidirectional measurement of causality from head motion to eye movement, does exist in the context of goal-directed sensorimotor tasks, as discussed in Section *Rationale of the Proposed UCM*.

According to its rationale, the proposed causality measure *UCM*_*Y →X*_ establishes a quantitative and normative definition, specially for the *specific head-eye coordination* involving head motions temporally preceding eye movements. *UCM*_*Y →X*_ > 0 offers a unidirectional cause-effect relationship. More specifically, *UCM*_*Y →X*_ > 0 represents that head motion is the cause and eye movement is the effect, *UCM*_*Y →X*_ < 0 indicates the causality from eye movement to head motion, and *UCM*_*Y →X*_ = 0 (practically *UCM*_*Y →X*_ approaches zero) means there is no unidirectional causality. In a word, based on the transfer entropies between head motion and eye movement, the causality measure *UCM*_*Y →X*_ obtains, objectively and quantitatively, a deeper understanding of the *specific head-eye coordination*.

### Significance Test

Measurement variance and estimation bias usually occur when obtaining transfer entropy, which is a common consideration [23]. Here we take a hypothesis testing approach [38] to combat this problem.

The standard statistical technique of hypothesis testing [23, 38, 39], due to its popular use in handling time series data, is performed for determining whether there exists a valid *UCM*_*Y →X*_ with a high confidence level. To do this, the null hypothesis *H*_0_ taken is that *UCM*_*Y →X*_ is small enough, that is, it means that *X* and *Y* do not influence each other. And *H*_1_ supports a causal-effect relationship between *X* and *Y*, unidirectionally. To verify or reject *H*_0_, surrogate time series 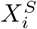 and 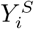 (*i* = 1, · · ·, *N*_*S*_) of the original *X* and *Y*, respectively, are used. For surrogate generation, random shuffle, which is simple yet effective, is utilized, because in this paper the history lengths of *X* and *Y* are both taken as 1, as usually used for the practical definition and computation of transfer entropy [23]. The unidirectional causality from the 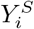 to 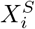, following (4), is obtained as follows:

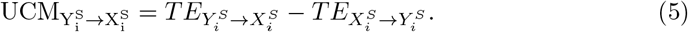

The significance level of *UCM*_*Y →X*_ is defined as:

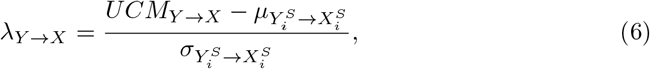

Where 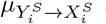 and 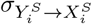 are the mean and standard deviation of 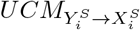 values, respectively. The probability to reject *H*_0_ can be obtained based on Chebyshev’s inequality, calculated as follows:

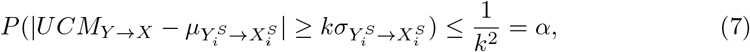

where 1 − *α* is the confidence level to reject *H*_0_ (and to accept *H*_1_), and parameter *k* is any positive real number. The number of the surrogates, which is related with the confidence level, is obtained as:

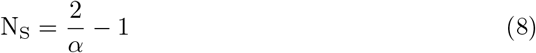

for a two-sided test.

In this paper, the parameter *k* used in (7) is taken as 6 resulting in a confidence level of 97.3% and, this is a high requirement satisfied in practice [39]. That is, if the significance level is bigger than 6 (*λ*_*Y →X*_ *>* 6), then, equivalently with a confidence level of more than 97.3%, there exists a unidirectional head-eye causality from head motion to eye movement (note this technique is called 6 − *Sigma* [38], some other techniques based on *p* − *value* approach to statistical significance testing [40] could be attempted for future). In fact, according to statistical test theory [39], it is important to know that a minimum confidence level, acceptable in practice, is 95.0% (here the corresponding significance level is 4.47). Notice that the significance and confidence levels play a same role in the hypothesis testing.

The *UCM*_*Y →X*_ and 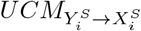 computations, highlighted by red and blue boxes respectively, are illustrated in Fig. 1. The example values of *UCM*_*Y →X*_ and 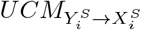 based on the gaze and head data by Participant 5 in Trial 3 in our psychophysical studies are also presented (here 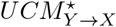and 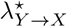 are especially used for emphasis, see all the results relevant to the unidirectional causality measure in Section *The Unidirectional Causality from Head Motion to Eye Movement*. Clearly, a big difference between 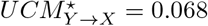 (with a very high confidence level of 99.3% and a very large significance level *λ*^*^ of 12.53) and 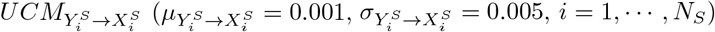, exists. Obviously, for the driving activity of this participant 5 in Trial 3, there appears significantly a unidirectional head-eye causality from head motion to eye movement in the goal-directed sensorimtor tasks.

**Fig 1.**
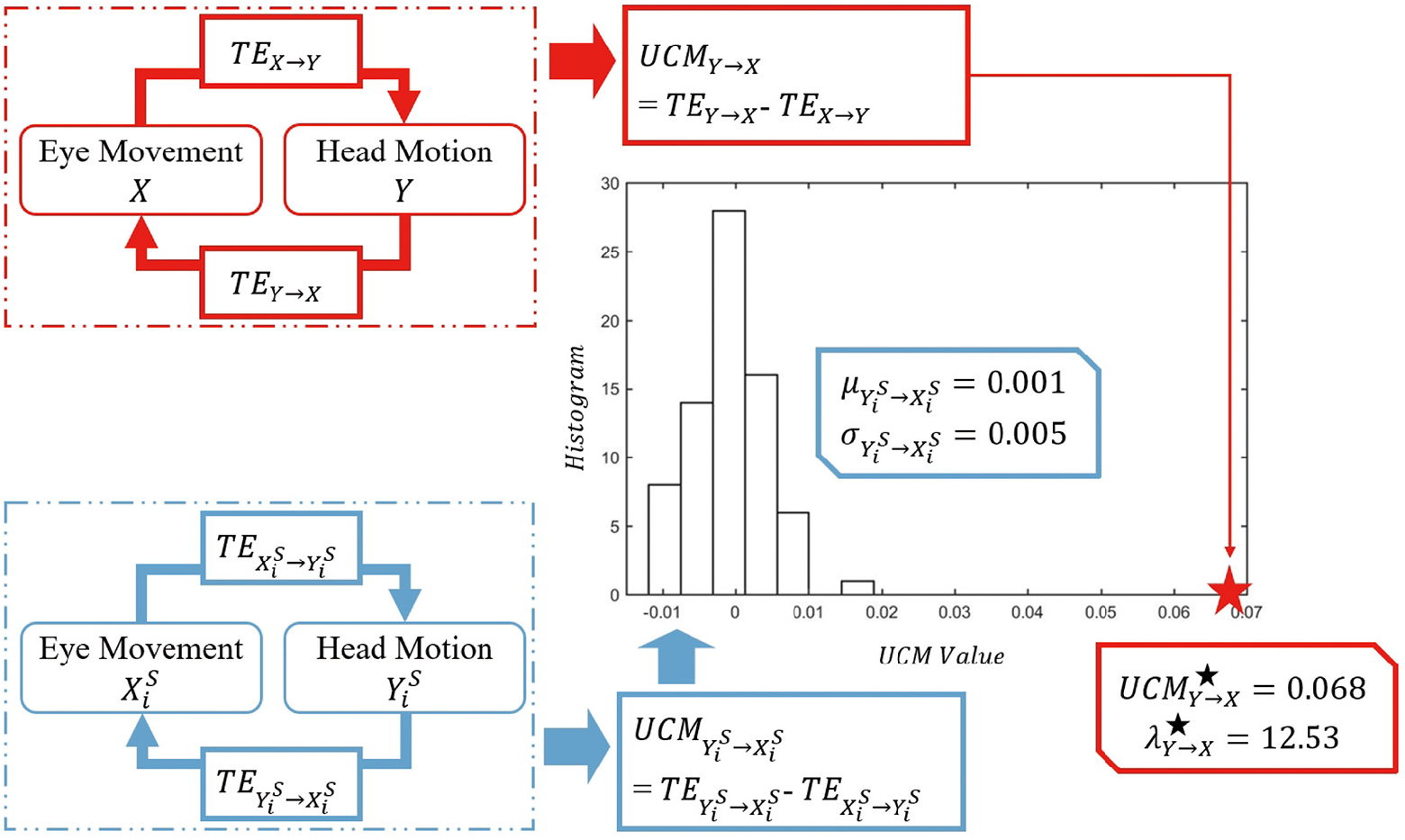
An illustration scheme for the computation of Unidirectional Causality Measure *UCM*_*Y →X*_.

It is noticed that the significance test for the computation of unidirectional causality measure described here is standard and general enough to be employed as well for checking the statistical significance of transfer entropy, as shown in Section *Transfer Entropies between Head Motion and Eye Movement*.

### Rationale of the Proposed UCM

The proposed unidirectional causality measure possesses the rationale due to its “physics” meaning and, to the verification by significance testing on itself.

During the procedure of goal-directed driving, the transfer entropy in the direction from head motion to eye movement *TE*_*Y →X*_, which is computationally verified by the significance test (as demonstrated in Section *Transfer Entropies between Head Motion and Eye Movement*) does exist. *TE*_*Y →X*_ provides the quantity of the basic content of causality in this direction. By contrast, the transfer entropy in the direction from eye movement to head motion *TE*_X→Y_, which is found with no statistical significance (also as demonstrated in Section *Transfer Entropies between Head Motion and Eye Movement*), actually means that the activity of visual scanning and visual-cognitive processing in goal-directed driving lacks the coordination dynamics with eye movements temporally preceding head motions. But very importantly, the *TE*_X→Y_ quantity itself does have a reasonable meaning. We believe that *TE*_X→Y_ can be intelligently considered as a value of the overall perturbations when obtaining the transfer entropy based causality in the direction from head motion to eye movement; as a result, *UCM*_*Y →X*_, which is *TE*_*Y →X*_ minus *TE*_X→Y_, takes an enhanced *TE*_*Y →X*_ and represents a unidirectional causality in this direction only. We call this is the “physics” meaning of the proposed causality measure *UCM*_*Y →X*_; moreover, according to the significance testing conducted on *UCM*_*Y →X*_, as demonstrated in Section *The Unidirectional Causality from Head Motion to Eye Movement*, the reasonableness of the “physically” based meaning of *UCM*_*Y →X*_ can be statistically verified. As a result, we can prove the rationale of the proposed unidirectional causality measure (*UCM*) and in this case, *UCM*_*Y →X*_ gives a determinant for the *specific head-eye coordination*.

## A Normalized Unidirectional Causality Measure (*NUCM*)

In a goal-directed driving scenario, the *specific head-eye coordination* corresponds to the state of visual scanning and visual-cognitive processing (correspondingly, the attentional state of drivers) [11–14], and meanwhile this state signifies the performance of sensorimotor tasks [21, 22]. As discussed in Section *A Unidirectional Causality Measure*, the unidirectional causality from head motion to eye movement in effect gives an quantitative estimation of the *specific head-eye coordination*. Therefore, we hypothesize that the unidirectional head-eye causality should work well as a proxy of the driving performance. This hypothesis will be verified by using the correlation analysis technique, which is a classic and popular tool for investigating a relationship between variables [41]. Because a proxy indicator of driving performance actually contributes to an objective and quantitative score for the sake of comparing performances, we propose a normalized unidirectional causality measure from head motion to eye movement, *NUCM*_*Y →X*_, for being quantitatively compatible with the driving performance, as follows:

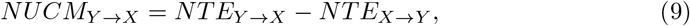

Where

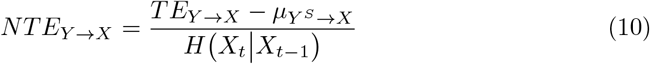

is a kind of normalized transfer entropy, whose definition is effective and popularly used [23]. Here *µ Y ^S^ → X* is the mean of the transfer entropies 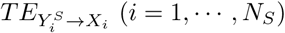 from surrogate head motion to original eye movement, the conditional entropy *H*(*X*_*t*_|*X*_*t*−1_) denotes the maximum of *TE*_Y→X_. *NTE*_*X*→*Y*_ can be obtained similarly,

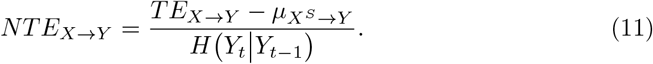

Note other normalization methods for transfer entropy and for the unidirectional causality measure could be done in future work [42, 43].

### Rationale of the Proposed NUCM

It is necessary to point out that the proposed normalized unidirectional causality measure *NUCM*_*Y →X*_ is rational because it keeps the rationale of *UCM*_*Y*→*X*_.

It is worthwhile to emphasize that *NUCM*_Y→X_ does keep the reasonable spirit of its “physically” based meaning, which is inherited from its original version *UCM*_*Y→X*_ (as discussed in Section *Rationale of the Proposed UCM*). That is, *NUCM*_*Y→X*_ characterizes a causality in the unidirectional direction only, from head motion to eye movement, indicating a determinant of the *specific head-eye coordination*. Then, the two components of *UCM*_*Y→X*_, *TE*_*Y→X*_ and *TE*_*X→Y*_, have separate and distinct functions, thus the normalization operations on these two transfer entropies should be handled separately for obtaining *NUCM*_*Y→X*_. *NUCM*_*Y→X*_ values range from −1 to 1 after normalization, providing a representation of the strength of the *specific head-eye coordination* that reflects the state of visual scanning and visual-cognitive processing and, the attentional state of drivers. As a result, *NUCM*_*Y→X*_ serves the practical purpose of performance comparison. In the procedure of performing a sensorimotor task, the bigger this strength of coordination is, the better the attentive/cognitive state of drivers becomes, the higher the task performance is; and *vice versa*. It is emphasized that the normalized unidirectional causality measure proposed can be easily and very well exploited in real applications such as safety driving, for providing an effective technique to monitor and obtain the attentional situations of drivers. And this exploitation offers a valuable and potential approach to behaviometrics for sensorimotor tasks in practical application scenarios.

## Experiment

### Virtual Reality Environment and Task

Driving, which is commonly considered as a goal-directed activity [7, 8], is taken as the sensorimotor task in our psychophysical experiments. Due to its repeatable usability, high safety and good performance, (head-worn) virtual reality technique has become a popular paradigm to study gaze shifts and sensorimotor tasks [9, 44–46]. Therefore, our study is done based on head-worn virtual reality.

In this paper, the virtual environment for the psychophysical studies utilizes a four-lane two-way suburban road consisting of straight sections, curves (4 left bends and 4 right bends with mean radii of curvature of 30 m) and 4 intersections, with common trees and buildings. In order to focus on the study of goal-directed activity in sensorimotor driving and on investigating quantitatively the *specific head-eye coordination* (with head motions temporally preceding eye movements), irrelevant visual distractors such as the sudden appearance of a running animal, which have been considered as ignored in the performing of goal-directed tasks [20] (and also this topic relevant to irrelevant visual distractors has been understood well in the research area [47]), are not included. Notice that, regardless of whether or not and how many irrelevant distractors are present, this *specific head-eye coordination* always and largely dominates in goal-directed sensorimotor tasks, as widely accepted in literature [1, 2, 7–9, 12] and as discussed in detail in Section *Introduction*.

In our study, a single driving task, which is to smoothly maintain the driving speed at 40 km/h, is used. The inverse of the average acceleration during driving is taken as the indicator of driving performance, as popularly done in literature [48]. That is, the larger the average acceleration is, the worse driving performance becomes, and *vice versa*.

Example illustrations of the virtual environment and of performing a driving task are presented in Fig. 2.

**Fig 2.**
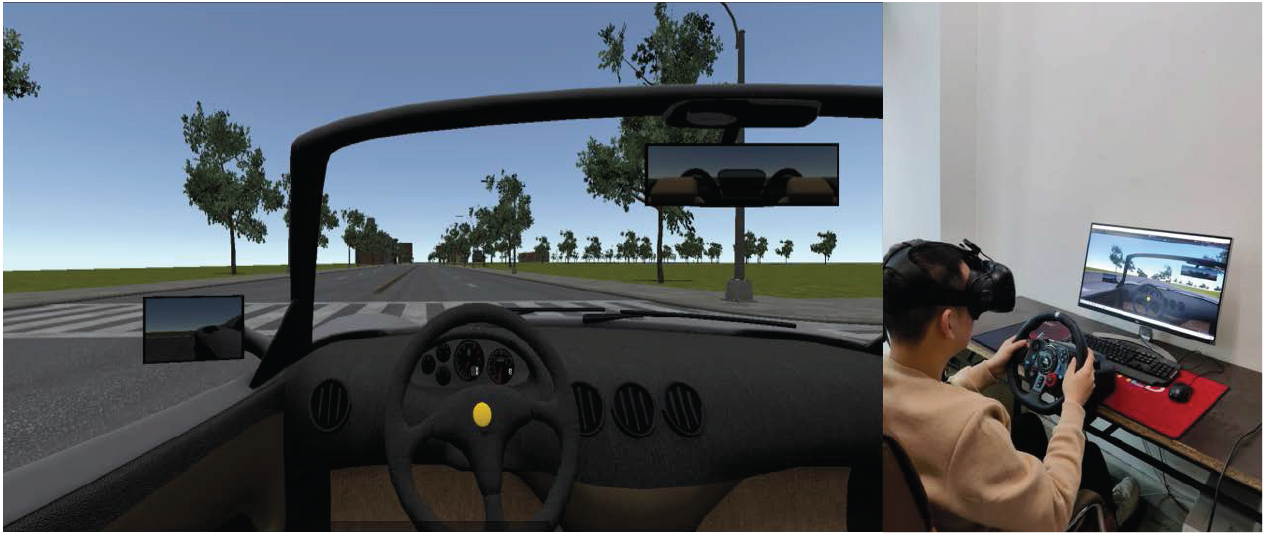
In the virtual environment (left), a participant is performing the driving task (right).

### Apparatus

The psychophysical experiments in this paper are conducted in a virtual reality environment through the display by an *HTC Vive* headset [49]. And there is a *7INVENSUN Instrument aGlass DKII* eye-tracking equipment [50] embedded in the headset. An illustration of the headset with the embedded eye tracker is given in Fig. 3.

**Fig 3.**
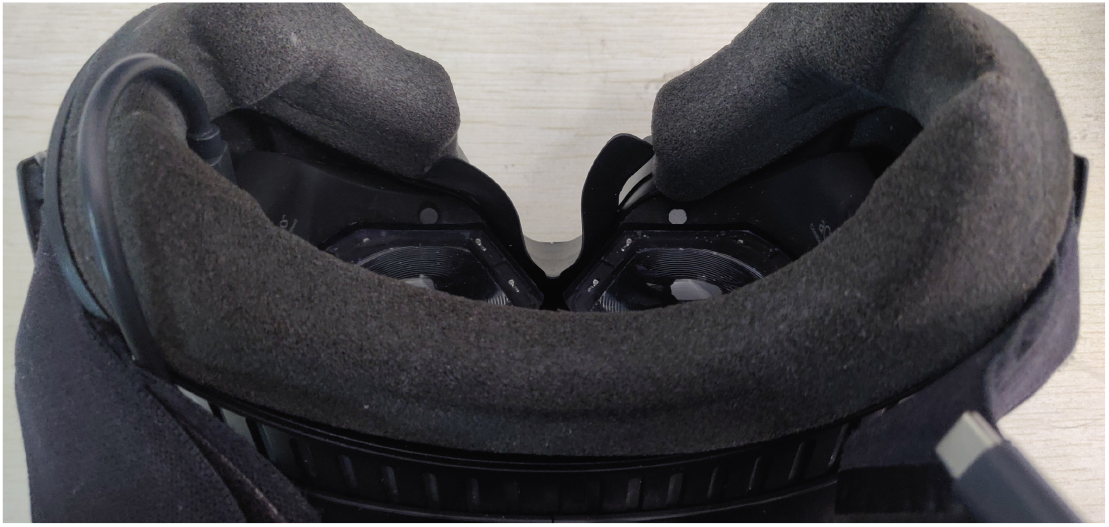
A *HTC Vive* headset and a *7INVENSUN Instrument aGlass DKII* eye tracker.

Eye movement (gaze point) and head motion (head pose) data are recorded in a frequency of 90 Hz by the eye-tracking equipment (gaze position’s accuracy is 0.5°) and by the headset respectively, both being captured as pitch and yaw (as usually done in the relevant filed [12]). Virtual driving is performed based on a *Logitech G29* steering wheel [51]. A desktop monitor is utilized to display the captured data and driving activities of participants in the procedure of experiment.

### Participants

Twelve people participated in the psychophysical study. Each participant took part in four independent test sessions to have a large enough sample size for our study (see details in Section *Procedure*). These participants, with normal color vision and normal/corrected-to-normal visual acuity, were recruited from students at one of the authors’ universities (7 male, 5 female; ages 22.9 ± 1.95). All of the participants hold their driver licenses for no less than one and a half years. None of the participants had any adverse reactions to the virtual environment utilized in this study. All participants provided written consent and were compensated with payment. This study was approved by the Ethics Committee of one of the authors’ universities under the title ‘Eye tracking based Quantitative Behavior Analysis in Virtual Driving’.

### Procedure

Each participant finished four test sessions, with an interval of one week between every two consecutive tests, based on the same task requirements and driving routes. In this study, a test session is represented as a trial. In total, there were 12 * 4 = 48 valid trials accomplished by the psychophysical experiments. Although this number of trials satisfies the large-sample condition in classical statistics [52], in the near future a larger sample size could be utilized for making our proposed causality measures get more possible contributions to practical behaviometrics applications.

Before each test, the purpose and procedure of psychological studies were introduced to the participants. For the sake of high-quality data recordings, (a) all participants completed a 9-point calibration procedure prior to the experiments; (b) the headset was adjusted and fastened to participants’ head; (c) sight and eye cameras were adjusted to prevent hair and eyelashes from obscuring; and (d) the seat was adjusted to a comfortable position in front of the steering wheel.

For each test session, first of all, conducting a 3-minute period of familiarization was introduced. Then, for a 3-min driving session, participants were instructed to comply with driving rules: driving smoothly at a speed of 40 km/h and following the formulated routes (trying to maintain close to the center line).

## Results and Discussions

### Temporal Sequences of Head Motion and Eye Movement Data

Example data for head motion and eye movement are plotted as a function of time, shown in Fig. 4. As usually done in the study of the coordination of head motion and eye movement [12, 14], the data of eye and head rotations in yaw are utilized in this paper. It is interesting to study further the pitch data, which will be as our future work. It is obvious that head motion and eye movement always exist during driving, even in the case of driving straight on a road (for instance, see the scenario during the first 1000 time units). Furthermore, the synchronized registration of the local extreme values of head motion and eye movement data indicates, to a certain extent, an overall correspondence between two kinds of data, clearly showing that the coordination of head and eyes does exist. Actually, we introduce an evaluation measure for the amount of coordination of head and eyes (*CoordAmount*), inspired by the widely used measure *PSNR* in the field of signal processing [53], as follows:

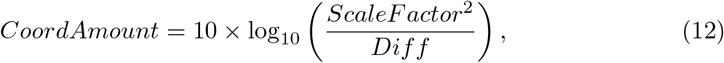

**Fig 4.**
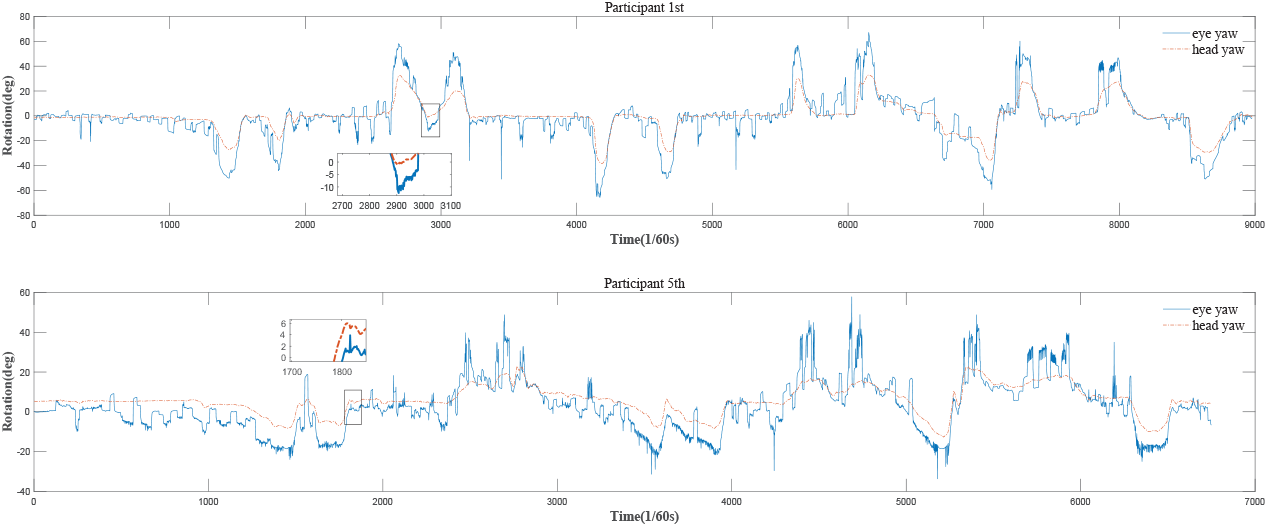
Examples of head and eye rotation data, from the fourth and third trials by two participants 1st and 5th, upper and lower rows, respectively. Basically, both head and eyes always rotate during driving, even during straight driving (see, for example, the scenario of the first 1000 time units). Particularly, head motion temporally precedes eye movement. For instance, in the boxes of the upper/lower rows, respectively, the head yaw starts to consistently increase/decrease from its local minimum/maximum earlier than the eye yaw, indicating a directional, complex and non-linear coordination between head and eyes. That is, the *specific head-eye coordination*, as a very natural human behavior, does come out even no irrelevant distractor presents. The coordination patterns by the two participants, which imply the inherent parts of their corresponding *specific head-eye coordination*, are largely different, as demonstrated with the statistics (mean (*M*) and standard deviation (*SD*)) of rotation data (*M*_*eye*_ = −1.35, *SD*_*eye*_ = 19.42, *M*_*head*_ = −0.46 and *SD*_*head*_ = 11.57, participant 1st in 4th trial; *M*_*eye*_ = 2.41, *SD*_*eye*_ = 12.90, *M*_*head*_ = 5.78 and *SD*_*head*_ = 7.06, participant 5th in 3rd trial). Correspondingly, the values of driving performances are 0.34 and 0.42 respectively, which are relatively diverse (the latter is 1.24 times as large as the former). The very distinct values of the proposed causality measures for these two participants are, respectively, 1.24 * 10^−2^ and 6.80 * 10^−2^ (*UCM*_*Y →X*_), together with −0.18 * 10^−2^ and 8.07 * 10^−2^ (*NUCM*_*Y →X*_). Obviously, the distinction between the two values of *UCM*_*Y →X*_ corresponds closely to the large difference between the two coordination patterns and, to the relatively large difference between the two performances. Further, the same situation applies to *NUCM*_*Y →X*_. What is stated here exactly means that, the *specific head-eye coordination* (and also the coordination pattern) can be quantitatively characterized by our new proposed causality measures, for indicating the attentive/cognitive state and driving performance. Finally and notably, the proposed *UCM*_*Y →X*_ and *NUCM*_*Y →X*_ can largely help discriminate between the performances of driving.

Where

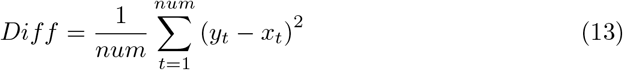

is the mean square difference (*Euclidean* distance) between gaze and head rotation data (*num* is the number of time units (*t*) considered). *ScaleFactor* = 360, is the maximal absolute difference between any two gaze and head data pair. *CoordAmount* quantifies a quality of matching two kinds of rotation data streams according to data values and shows the synergy of both rotation streams, providing a normalized measurement of the amount of synergistic coordination of head and eyes. The more amount of coordination exists, the higher the *CoordAmount* becomes, and *vice versa*. The *CoordAmount* values, which are 31.45 dB and 33.09 dB for participants 1st (fourth trial) and 5th (third trial) respectively, are relatively high and, this verifies the existence of the coordination of head and eyes. Note these two amounts of coordination for the two trials are close.

However in fact, *CoordAmount* does not provide an answer to the definition of the coordination of head and eyes itself. A major feature of coordination interactions between head motion and eye movement is that the head moves earlier than eyes, as specially illustrated by the boxes in Fig. 4. In the box of the upper row, the head yaw starts to consistently increase from its local minimum earlier than the eye yaw, and in the box of the lower row, the head yaw starts to consistently decrease from its local maximum earlier than the eye yaw. This indicates a temporally directional and non-linear coordination relationship between head and eyes, and namely the so-called *specific head-eye coordination*. Actually this specific coordination is usually and always involved in the driving procedure for the fulfillment of goal-directed sensorimotor tasks, irrespectively of whether or not and how many irrelevant distractors present, as suggested in the literature [11–14]. Even no irrelevant visual distractor appears, the *specific head-eye coordination* does come out because it is a natural and essential part of human behavior [7, 9, 19]. In addition, detail examination of the head motion and eye movement data finds that the coordination patterns by the two participants are largely distinct. For example, in Fig. 4 the mean (*M*) and standard deviation (*SD*) values for these data are as follows: *M*_*eye*_ = −1.35, *SD*_*eye*_ = 19.42, *M*_head_ = −0.46 and *SD*_*head*_ = 11.57, participant 1st in the fourth trial; *M*_*eye*_ = 2.41, *SD*_*eye*_ = 12.90, *M*_*head*_ = 5.78 and *SD*_*head*_ = 7.06, participant 5th in the third trial. Indeed, the coordination pattern of the head and eyes implies an inherent part of the *specific head-eye coordination*, as mentioned by literature [12]. Furthermore, as found by the relevant literature [12, 54], a large distinction with coordination patterns corresponds to big different states of attention in sensorimotor activities and thus, diverse driving performances. For the example demonstrated in Fig. 4, the corresponding performance values are relatively diverse, 0.34 and 0.42 for the two participants respectively. The latter is 1.24 times as large as the former. Therefore, it would be curious to know how the *specific head-eye coordination* is closely related with driving activity and performance. Actually, all of this means that it is not easily clear how to obtain an quantitative definition of the coordination of head and eyes, based directly on the use of head motion and eye movement data and/or of their statistics (such as mean and standard deviation), for the context of evaluation of driving performance. In particular, it is challenging to define an quantitative measure for the *specific head-eye coordination*, due to the fact that this special coordination involves the directional, complex and non-linear problem of probabilistic prediction, as indicated in the relevant literature [12] (also, as demonstrated by the boxes in Fig. 4 and, by the proof from our results in Section *Transfer Entropies between Head Motion and Eye Movement*). Fortunately, this challenge has been overcome by the proposed causality measures (*UCM* and *NUCM*) based on transfer entropy. And as a matter of fact, the new causality measures provide an indicator of the coordination (pattern) of head and eyes, also essentially our proposal of causality philosophy reveals a determinant for the *specific head-eye coordination* under consideration (as to be demonstrated in detail in Sections *The Unidirectional Causality from Head Motion to Eye Movement & The Normalized Unidirectional Causality from Head Motion to Eye Movement*).

### Transfer Entropies between Head Motion and Eye Movement

All the values of transfer entropies are listed in Table 1. A significant difference between two transfer entropies *TE*_*Y →X*_ and *TE*_*X→Y*_ is revealed by one way analysis of variance

**Table 1.** Values of Transfer Entropies *TE*_*X→Y*_ and *TE*_*Y →X*_ (*10^−2^)

| Participant | Trial 1 |  | Trial 2 |  | Trial 3 |  | Trial 4 |  |
| --- | --- | --- | --- | --- | --- | --- | --- | --- |
| | $X \rightarrow Y$ | $Y \rightarrow X$ | $X \rightarrow Y$ | $Y \rightarrow X$ | $X \rightarrow Y$ | $Y \rightarrow X$ | $X \rightarrow Y$ | $Y \rightarrow X$ |
| 1 | 2.19 | 2.98 | 1.99 | 2.77 | 1.68 | 1.85 | 1.75 | 2.99 |
| 2 | 1.82 | 3.03 | 2.60 | 6.93 | 1.64 | 3.52 | 2.00 | 4.76 |
| 3 | 2.25 | 3.68 | 1.97 | 2.58 | 2.42 | 2.72 | 2.66 | 4.48 |
| 4 | 0.79 | 1.97 | 1.25 | 2.23 | 1.31 | 3.16 | 1.17 | 3.86 |
| 5 | 1.82 | 4.96 | 1.13 | 5.87 | 0.86 | 7.66 | 1.59 | 3.84 |
| 6 | 1.83 | 5.38 | 4.28 | 8.71 | 2.06 | 5.09 | 1.99 | 3.96 |
| 7 | 1.45 | 4.36 | 2.99 | 6.41 | 2.26 | 6.60 | 1.09 | 4.74 |
| 8 | 1.39 | 2.31 | 1.49 | 3.51 | 3.00 | 6.86 | 2.31 | 3.88 |
| 9 | 1.76 | 5.09 | 2.17 | 6.50 | 1.85 | 5.60 | 2.16 | 3.79 |
| 10 | 1.12 | 1.80 | 1.23 | 4.30 | 1.08 | 2.99 | 1.86 | 4.95 |
| 11 | 2.27 | 5.02 | 1.22 | 2.43 | 2.09 | 3.98 | 1.10 | 2.80 |
| 12 | 1.32 | 3.58 | 2.34 | 7.87 | 2.81 | 8.99 | 2.36 | 9.37 |

(*ANOVA*) (*F* (1, 94) = 80.25, *p <* 0.05), as illustrated in Fig. 5. Obviously, the transfer entropy in the direction from head motion to eye movement, *TE*_*Y →X*_, is much bigger than that in the reverse direction, *TE*_*X→Y*_, with the averages of the former and latter, 3.8 *10^−2^ and 1.9 * 10^−2^, respectively. That is, *TE*_*Y →X*_ is twice as big as *TE*_*X→Y*_ for the experimentation data in this paper.

**Fig 5.**
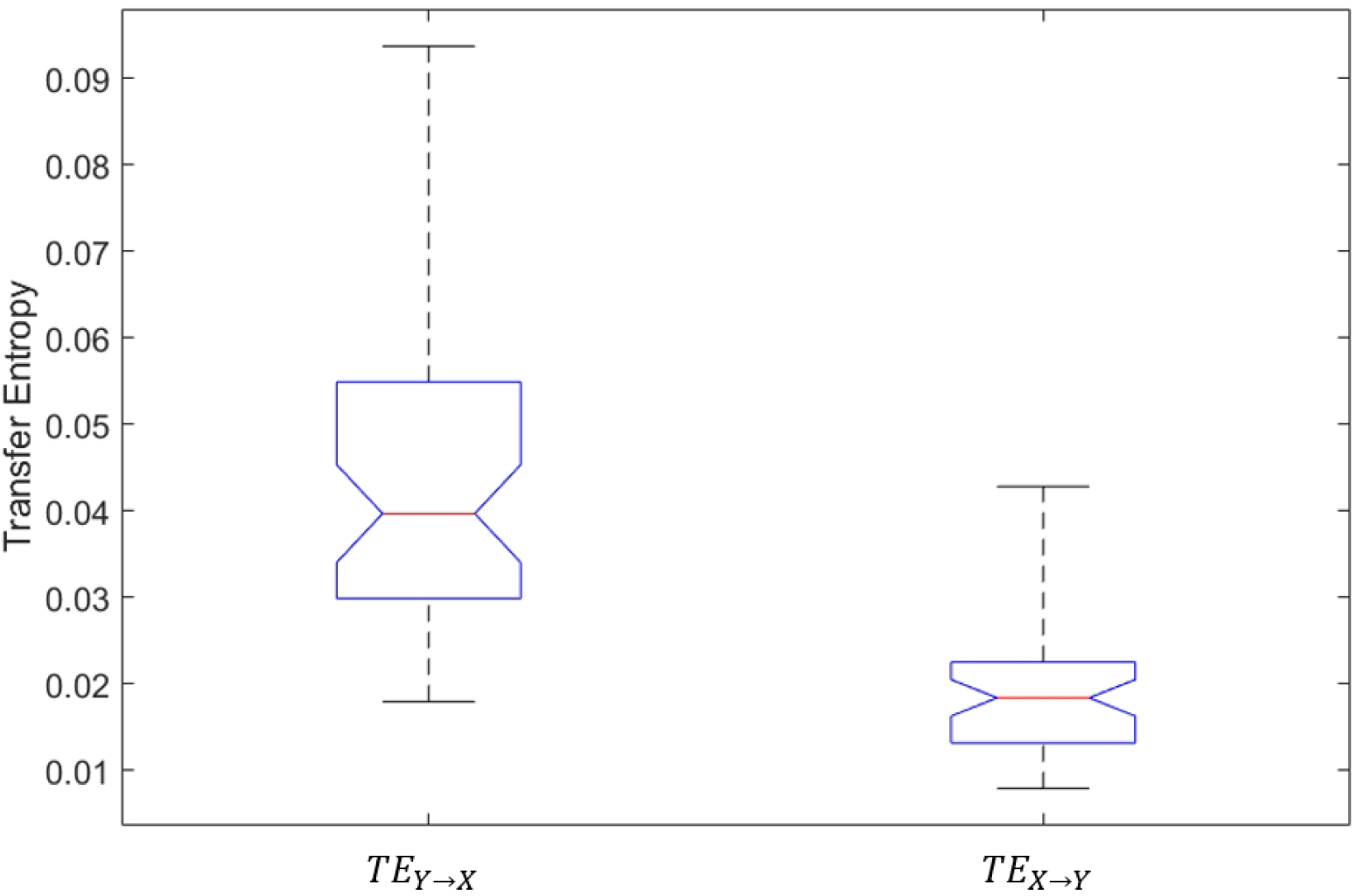
Transfer entropies between head motion and eye movement.

Further, a statistical significance testing, which is completely similar to what has been described in Section *Significance Test* and also in literature [23], is used for checking the statistical confidence levels of *TE*_*Y →X*_ and *TE*_*X→Y*_, entirely separately. It is observed that the significance and confidence levels for *TE*_*Y →X*_ are 4.49 and 95.0%, respectively. In contrast, the two corresponding values for *TE*_*X→Y*_ are 1.46 and 53.4%, respectively. This means that *TE*_*Y →X*_ and *TE*_*X→Y*_ are statistically acceptable and unacceptable, respectively. Therefore, the coordination of head and eyes with head motions temporally preceding eye movements, indicated by the information flow from head motion to eye movement *TE*_*Y →X*_, is validly and reliably evidenced with a high probabilistic certainty; but on the contrary, the transfer of information from eye movement to head motion (indicated by *TE*_*X→Y*_, namely the directional coordination dynamics with eye movements temporally preceding head motions) cannot be identified in this goal-directed sensorimotor task.

The results in the above two paragraphs for transfer entropies, quantitatively, validate and express that the coordination between head and eyes in the visual scanning for performing sensorimotor tasks always exists [10–14]. Particularly, the *specific head-eye coordination* involving head motions temporally preceding eye movements is mainly occupied in a goal-directed and task-driven driving task, irrespectively of whether or not and how many irrelevant visual distractors appear [1, 2, 7–9]. Here, notably, even no irrelevant visual distractor presents, the *specific head-eye coordination* does exist because it represents a natural essence of human behavior [7, 9, 19]. It is highlighted that the only validity of the unidirectional transfer entropy from head motion to eye movement witnesses the non-linearity of the *specific head-eye coordination*, as demonstrated with the boxes in Fig. 4. And also, the idea of characterizing an information flow by transfer entropy matches the essence of this directional coordination.

### The Unidirectional Causality from Head Motion to Eye Movement

The unidirectional head-eye causality *UCM*_*Y →X*_ results (as provided in Table 2) are obtained with high significance levels (*λ*_*Y →X*_), which are presented in Table 3. Almost all the *λ*_*Y →X*_ values are larger than 6, that is, the corresponding confidence levels are more than 97.3%. There are only two exceptional evaluations of *λ*_*Y →X*_, 5.7 and 5.5, marked with boxes (Table 3), are slightly lower than 6. Even here, the corresponding confidence levels are 96.9% and 96.7% respectively and, this is acceptable in statistics for practical use [55]. The strict positive *UCM*_*Y →X*_ (Table 2) reveals that there indeed exists a unidirectional causality from head motion to eye movement (with high confidence), in the procedure of performing goal-directed sensorimotor tasks. Notably, the creativity of our work, considering normatively the *specific head-eye coordination* as defined as a unidirectional causality measure in a calculable fashion, is strongly verified by this discovery. And in the meanwhile, our causality discovery here objectively and quantitatively proves the observation that, during performing sensorimotor tasks, visual scanning is mainly done through the unidirectional and non-linear coordination of head and eyes involving head motions temporally preceding eye movements, even though no irrelevant visual distractor appears because this coordination is a natural behavior [7, 9, 19]. As a matter of fact, the existence of the proposed unidirectional causality here is an enhanced proof for the identification of the *specific head-eye coordination* and, for the presence of the unidirectional transfer entropy from head motion to eye movement.

**Table 2.** Values of Unidirectional Causality Measure *UCM*_*Y →X*_ (*10^−2^)

| Participant | Trial 1 | Trial 2 | Trial 3 | Trial 4 |
| --- | --- | --- | --- | --- |
| 1 | 0.79 | 0.78 | 0.17 | 1.24 |
| 2 | 1.21 | 4.33 | 1.88 | 2.76 |
| 3 | 1.43 | 0.61 | 0.30 | 1.82 |
| 4 | 1.18 | 0.98 | 1.85 | 2.70 |
| 5 | 3.14 | 4.73 | 6.80 | 2.25 |
| 6 | 3.55 | 4.43 | 3.02 | 1.97 |
| 7 | 2.91 | 3.42 | 4.33 | 3.65 |
| 8 | 0.91 | 2.02 | 3.86 | 1.57 |
| 9 | 3.33 | 4.33 | 3.75 | 1.62 |
| 10 | 0.68 | 3.06 | 1.91 | 3.09 |
| 11 | 2.76 | 1.21 | 1.89 | 1.70 |
| 12 | 2.26 | 5.53 | 6.18 | 7.01 |

**Table 3.** Significance Levels *λ*_*Y →X*_ for *NUCM*_*Y →X*_

| Participant | Trial 1 | Trial 2 | Trial 3 | Trial 4 |
| --- | --- | --- | --- | --- |
| 1 | 8.89 | 9.18 | 15.25 | 10.23 |
| 2 | 17.33 | 18.08 | 7.83 | 5.68 |
| 3 | 17.01 | 5.48 | 11.20 | 11.64 |
| 4 | 11.94 | 15.03 | 12.25 | 16.67 |
| 5 | 16.02 | 12.18 | 12.53 | 8.02 |
| 6 | 15.86 | 20.18 | 21.13 | 17.15 |
| 7 | 17.28 | 19.36 | 26.34 | 22.89 |
| 8 | 7.86 | 13.92 | 13.58 | 10.74 |
| 9 | 16.40 | 17.84 | 17.37 | 13.94 |
| 10 | 14.32 | 16.86 | 13.14 | 14.47 |
| 11 | 19.35 | 12.71 | 18.87 | 19.43 |
| 12 | 14.67 | 12.52 | 16.69 | 15.06 |

Besides, due to its rationale discussed in detail in Section *Rationale of the Proposed UCM*, the quantitatively unidirectional head-eye causality measure (*UCM*_*Y →X*_) gives an objective and explainable determinant of the *specific head-eye coordination*, which is important to be emphasized. For an illustrative example (see Fig. 4 and the corresponding descriptions, Section *Temporal Sequences of Head Motion and Eye Movement Data*), the *UCM*_*Y →X*_ values achieved from the fourth and third trials by the participants 1st and 5th are largely distinct, with 1.24 * 10^−2^ and 6.8 * 0 10^−2^, respectively. The latter is 5.4 times as big as the former. The substantial distinction here is in a high correspondence with the large difference between the two coordination patterns of head and eyes by these two participants but, in a big contrast to the closeness between the two corresponding values of amount of coordination (*CoordAmount*). Considering that the coordination pattern of head and eyes inherently contain an implication of the *specific head-eye coordination*, as discussed in Section *Temporal Sequences of Head Motion and Eye Movement Data*(alshythe illustration here evidences that the unidirectional causality from head motion to eye movement gives an underlying determinant of this coordination.

Despite the fact that the history length used in the time series data of this paper is simply taken as 1, as usually used in literature [23], and has shown satisfactory results, we would try other possible options of history length in future work. It is also worthwhile noting that *UCM*_*Y →X*_ is obtained by hypothesis test based on the application of surrogate time series [39], and a random shuffle is performed due to the present use of history length for obtaining the surrogate being utilized. As for other possible methods of generating surrogate time series, iterative amplitude adjusted Fourier transform (*iAAFT*) [56] could be suitable. Considering that hypothesis test is a necessary computational component to estimate causality measures, hypothesis test by *e-values* [57] would be as a more effective solution for future.

### The Normalized Unidirectional Causality from Head Motion to Eye Movement

Results of the normalized unidirectional head-eye causality (*NUCM*_*Y→X*_) and the corresponding driving performance (the inverse of the average acceleration, denoted by 1*/AvgAcc*) are listed in Table 4. As a concrete instance, depicted in Fig. 4 together with the corresponding descriptions in Section *Temporal Sequences of Head Motion and Eye Movement Data*, the two very different *NUCM*_*Y→X*_ values by the participants 1st and 5th (in the trials 4th and 3rd) are −0.18 * 10^−2^ and 8.07 * 10^−2^, respectively. The large difference between these two values of *NUCM*_*Y→X*_ corresponds closely to the big difference between the two coordination patterns of head and eyes by these two participants and meanwhile, contrasts sharply with the closeness of the two corresponding *CoordAmount* values. Importantly, this clearly reveals that the proposed *NUCM*_*Y→X*_, which is as an indicating pointer to well characterize the *specific head-eye coordination* (and also the coordination pattern), is largely related with driving activity and performance. More importantly, *NUCM*_*Y→X*_ is very and even enhanced discriminating to differentiate the distinct driving activities of the two participants under consideration (correspondingly, the two relatively diverse values of driving performances are 0.34 and 0.42 respectively, with the latter is 1.24 times as big as the former). In fact, a significant correlation (*p <* 0.05) between the new normalized causality measure and driving performance, based on all the head and gaze data in 48 trials, is obtained by three correlation analyses [41], with Pearson linear correlation coefficient (*PLCC*), Kendall rank order correlation coefficient (*KROCC*) and Spearman rank order correlation coefficient (*SROCC*) of 0.32, 0.27 and 0.41 respectively (Table 5). These correlation coefficient values, which range from moderate to relatively strong, definitely indicate a statistically significant relationship between our proposal of normalized causality measure and driving performance, as popularly recognized in the literature [58]. By contrast, the measurements by the compared techniques (Table 5) cannot show an acceptable association with the performance of virtual driving (*p >* 0.05).

**Table 4.**
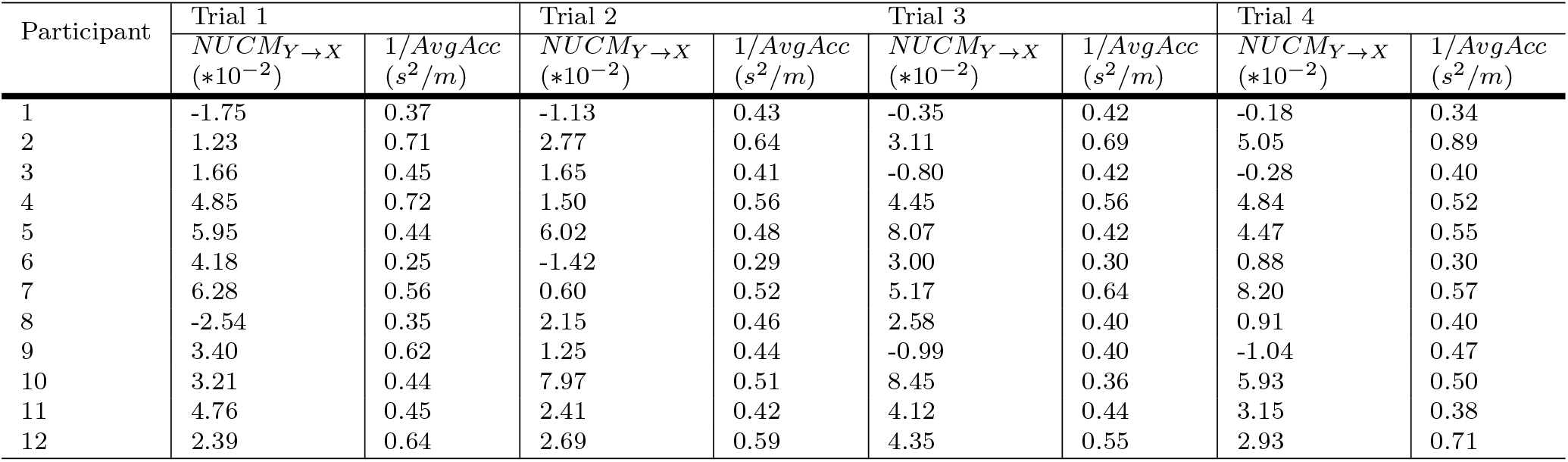
Values of Normalized Unidirectional Causality Measure *NUCM*_*Y→X*_ (*10^−2^) and driving performance (1*/AvgAcc*)

**Table 5.** Correlation Analysis between Measures and Driving Performance

| Methods | *PLCC*, $p$ - value | *KROCC*, $p$ - value | *SROCC*, $p$ - value |
| --- | --- | --- | --- |
| ***NUCM*<sub>$Y \rightarrow X$</sub>** | **0.32, $p < 0.05$** | **0.27, $p < 0.05$** | **0.41, $p < 0.05$** |
| *TGTE* | 0.19, $p > 0.05$ | 0.19, $p > 0.05$ | 0.26, $p > 0.05$ |
| *GTE* | 0.07, $p > 0.05$ | 0, $p > 0.05$ | -0.01, $p > 0.05$ |
| *SGE* | -0.07, $p > 0.05$ | -0.09, $p > 0.05$ | -0.15, $p > 0.05$ |
| *EoFS* | 0.01, $p > 0.05$ | 0, $p > 0.05$ | -0.03, $p > 0.05$ |
| *Entropy rate* | -0.06, $p > 0.05$ | -0.01, $p > 0.05$ | -0.02, $p > 0.05$ |
| *Fixation rate* | -0.24, $p > 0.05$ | -0.17, $p > 0.05$ | -0.24, $p > 0.05$ |
| *Saccade amplitude* | 0.25, $p > 0.05$ | 0.11, $p > 0.05$ | 0.09, $p > 0.05$ |

In brief, all the results demonstrated just above suggest that, in the procedure of performing goal-directed sensorimotor tasks, the proposed normalized unidirectional causality from head motion to eye movement, *NUCM*_*Y →X*_, offers a good indicator of task performance, providing a practical link with application of behaviometrics. Very notably, what is revealed is the first finding to clearly and quantitatively establish a connection between the *specific head-eye coordination* based causality measure and driving performance. Additionally, it is worthwhile to emphasize that the good achievement of the new normalized causality measure results from a strong management of the *specific head-eye coordination* as a quantitatively normative definition of causality. Also, the reason is that this specific coordination is usually and always involved in the visual scanning during goal-directed driving [11–14]. In particular, no matter whether distracting stimuli occur and no matter how many distractors appear, the *specific head-eye coordination* does exist [1, 2, 7–9]. It is noted that, even though no irrelevant distractor appears, this directional head-eye coordination still does exist because it is an intrinsic part of human behavior [7, 9, 19] and, it also indicates the cognitive state for whether or not and how focusing attention on visual scanning and driving [12–14, 21, 22]. All in all, due to its stable existence, the *specific head-eye coordination* can be well and further exploited to obtain *NUCM*_*Y →X*_, for the purpose of evaluating the attentive/cognitive state and driving performance (as to be discussed in the following paragraph).

Notice that the proposed normalized unidirectional causality from head motion to eye movement, *NUCM*_*Y→X*_, explicitly defines a directional measure based on the flow of information (transfer entropy), which depicts exactly the essence of the *specific head-eye coordination* involving head motions temporally preceding eye movements. It is important to note this essence manifests a natural and fundamental characteristic of goal-directed activity in performing a sensorimotor task (such as walking and driving). Actually, such a characteristic asserts a systematically temporal sequencing of activities achieved by different body components (including head, eyes, torso, hands, legs and so on) for indicating the attention and cognitive state of performers [8, 9, 12] and, this directional sequencing does relate to the performance of sensorimotor tasks [12, 54]. In our work, a typical sequencing completed by the coordination of head and eyes is specially considered. That is to say, the naturally happening *specific head-eye coordination*, which is driven by the performers’ knowledge and experience based on their goals [7, 9, 19], takes an indicator of the cognitive state for whether or not and how concentrating attention on visual scanning and driving [12–14, 21, 22]. Consequently, the proposed normalized causality measure (*NUCM*), which is of course in correlation with the attentive/cognitive state, can play a role of performance evaluation for driving.

It is crucial to emphasize that the definition of *NUCM* is essentially and totally different from that of the amount of coordination of head and eyes (*CoordAmount*, as introduced and discussed in Section *Temporal Sequences of Head Motion and Eye Movement Data*). *UCM* (in bits) represents a unidirectional causality measure by characterizing the directional information flow stemming from the time series of head motion to those of eye movement. *NUCM* is a normalized version of *UCM*, keeping the unidirectional characteristic of *UCM*. *CoordAmount* (in dB) gives a normalized version of the mean square *Euclidean* distance between two sequence signals of head motion and eye movement, representing the quantitative amount of coordination from the perspective of data-matching effectiveness. CoordAmount does not take into account the directional relationship of two sequences.

To go further, our proposal has achieved soundly an objective evaluation of the activity of visual scanning and visual-cognitive processing and, this achievement is, first of all, obtained through the exploitation of the transfer entropy based complexity measurement on unidirectional and non-linear head-eye relationship, involved in the specific coordination dynamics with head motions temporally preceding eye movements. Additionally, entropy based complexity measures, as versatile and powerful tools, have been verified and accepted for assessing the behavior (efficiency and performance, for example) of sophisticated systems [59]. Note that the results by the entropy based approaches under comparison (Table 5) cannot satisfactorily give an approximation to the performance of driving, but all these measures are entropy and complexity. Thus, how this happens is an amazing question. The answer is as follows. *EoFS* [36] (including entropy rate [35]) and *SGE* [37] give the definition of complexity for spatially based patterns of fixation sequence and of fixation, respectively. *GTE* [37] and *TGTE* [14] provide the complexity for, spatially and temporally respectively, based patterns of fixation transition. It is obvious that the compared entropy based measures just care about the information on fixation. In this paper, fixations are aligned with *AOI* s that have a one-one correspondence to 3D objects in dynamic virtual reality environments, as has been suggested in literature [8, 45]. Our proposal furnishes the complexity for causally and non-linearly based patterns of ordered pair between past head motion and current eye movement. Simply put, the proposal by us notably enables, rather than the space-time consideration of fixation relevant information only, the mastering of non-linear and unidirectional coordination dynamics from head motion to eye movement. Our methodology promotes a better consideration for human environment interactions. That is, what we have achieved is, more adequately, in agreement with the essential and goal-directed mechanism of visual scanning and visual-cognitive processing in sensorimotor driving.

Finally, eye tracking indices (such as fixation rate, which is the number of fixations in a unit time interval, and saccade amplitude), which are simple statistics on fixation/saccade, do not have correlation with driving performance (Table 5), though they are necessary for understanding visual scanning and visual-cognitive processing, especially aimed at particular application scenarios [60].

### General Discussions

Our paper aimed to investigate a human behavior, the coordination of the head and eyes, involved in human-environment interactions. We have attempted to address a quantitative and normative measure of the *specific head-eye coordination* in the goal-directed sensorimotor activities. We have successfully answered the following three questions: (a) Does the *specific head-eye coordination* essentially indicate a conceptual idea of causality?; (b) Can we normatively provide an objective description of the *specific head-eye coordination*?; and (c) Can we clearly find a quantitative relationship between the *specific head-eye coordination* and the performance of a sensorimotor task?

Importantly, the causality philosophy proposed in this paper represents a profound synthesis of ideas from the *specific head-eye coordination* in the goal-directed sensorimotor activity and the formal concept of Wiener&Granger causality. Our results on the unidirectional head-eye causality measures, proposed for visual-cognitive processing during processing goal-directed sensorimotor tasks, are particularly and quantitatively, in line with the combination of three relevant research lines. The first claims that the coordination of head and eyes is necessarily included [7, 11]. The second informs that the *specific head-eye coordination* with head motions temporally preceding eye movements, as a goal-directed mechanism, underpins the activity of visual scanning and visual-cognitive processing [11–14]. In addition, this specific coordination is usually and always involved whether or not there exist irrelevant visual distractors [1, 2, 7–9]. The third suggests that the state of visual scanning and visual-cognitive processing, namely the state of attention, is closely relevant to task performance [12–14, 21, 22]. To the best of our knowledge, this work is an original finding to clearly establish, in an objective and calculable manner, an effective linkage between the *specific head-eye coordination* and driving performance, providing a definition of efficiency measure for visual scanning and/or visual-cognitive processing, grounded on the exploitation of the entropy-based complexity for unidirectional relationship patterns of head-eye coordination. Simply, the proposed causality philosophy makes good use of the concept of information flow for giving a quantitative characterization of the *specific head-eye coordination*, indicating numerically the degree of goal-directed modulation and also the state of attention during driving.

With the development of the latest underpinning research of the cross-modal interplay between different body components (such as the synchronization of neural oscillations) [61], other kinds of comprehensive and systematic interplay between brain, visceral organs, head, eyes, and other body components are all future avenues of study. This is because they are all employed in the visual-cognitive processing for sensorimotor driving [25]. In this case, the proposed causality philosophy can be “generally” exploited. Even within the eye-tracking field, different types of measures of human behavior (such as pupil dilation and eyeblink rate) should logically and theoretically have complex relationships for the performing of cognitive tasks [62] and, the proposed causality philosophy can be taken advantage of. The generalization of the proposed causality philosophy, we do believe, may apply to all the possible causal-effect relationships, cross-modally and/or within-modally, involved in cognitive processing during (sensorimotor) behavior of human.

Currently, for the two proposed causality measures, we consider more from the probabilistic perspective of prediction and complexity, based on the utilization of an information flow tool, transfer entropy, holding the idea of prediction by Wiener&Granger causality [27, 28]. Although it is not so clear whether Pearl’s causality driven by the idea of intervention [31] works well in the application context without intervention, as has been recognized in the literature [63], a combination of the ideas of information flow (transfer entropy) and Pearl’s intervention would be approached, constituting the authors’ future work.

Interestingly and importantly, the goal-directed prediction has been accepted as a very recent research perspective on visually based perception and cognition [2]. As a typical goal-directed prediction, the activity of visual scanning and visual-cognitive processing in the procedure of performing sensorimotor tasks is produced based on the dialogue and bridge between top-down modulations and bottom-up stimuli [26, 61, 64]. The management of dialogue and bridge is factually witnessed and quantitatively obtained by the complexity measurement (transfer entropy) for causal and non-linear patterns of the *specific head-eye coordination*. Thanks to the outstanding ability of (transfer) entropy in the sense of *Shannon* to characterize the complexity of a sophisticated system, the two proposed cause-effect measures, as an objective indicator of goal-directed prediction, provide a natural, powerful, and original assessment of the performance for the fulfillment of sensorimotor tasks. Indeed, information physics [65] could be an emerging paradigm to help the further investigation, from the perspective of the complexity of a dynamic system. Another interesting consideration is to exploit the local information transfer [66] to study the dynamics of complexity system in our context.

In this paper, the inverse of the average acceleration, as popularly used in relevant literature [48], is taken as the driving performance, and this is in accordance with the task goal, namely driving smoothly and keeping driving speed constant. It is obvious that the fulfillment of this kind of driving essentially helps observe the goal-directed essence of sensorimotor activity. And meanwhile, the virtual environment is specially designed to use the common suburb road typically with straight sections and bends. All in all, the proposed causality measures can be investigated and evaluated in a focused way.

Last but also important, many practical applications, such as the estimation of human behavior in driving [12, 14, 67], sports [68] and health/illness [69], can take advantage of our proposed causality measures for the discovery and application of behaviometrics, as has been emphasized in Section *A Normalized Unidirectional Causality Measure*.

## Conclusion and Future Works

In this paper, major progress has been achieved in deeply understanding the *specific head-eye coordination*, with head motions temporally preceding eye movements, as a fundamental human behavior in visual scanning and visual-cognitive processing. Two new quantitative measures, namely the unidirectional causality measure and its normalized version, have been presented to objectively depict the *specific head-eye coordination*. Based on psychophysical studies conducted by using virtual reality, we have distinctly discovered that, during virtual driving, the two proposed causality measures behave very well as indicators for the cause-effect link between head and gaze movements, and for driving performance, respectively. Our proposed causality philosophy is simple but effective for obtaining the attention and cognitive state of task performers. To sum up, the two new causality measures have illuminated a quantitative and complete path, connecting well the three fundamental factors (namely, mechanism, attentional state, and task performance) involved in visual scanning and visual-cognitive processing during sensorimotor tasks. Our discovery undoubtedly offers a large potential for behaviometrics applications in the context of sensorimotor tasks.

In the future, quite a few considerations have been suggested for breaking the limitations mentioned in the above texts. The coordination relationship and temporal alignment are very fundamental in human behavior for diverse natural tasks [9, 10], and as a result, the proposed philosophy of a unidirectional causality based on transfer entropy provides a quantitative and universal potential for the feasibly operable measurement of these relationship and alignment. For instance, the complex coordination dynamics among head, eyes, torso, hands, and legs [9, 10] will be extensively studied, for the purpose of wider application and generalization of the proposed causality philosophy. For another instance, the unidirectional flow of information from eye movement to head motion, which results from stimulus-driven shifts of attention [70, 71], will be investigated for its quantification by the exploitation of transfer entropy. Particularly, in this case, the complex task with multiple task-relevant distractors and targets competing for gaze allocation [64] would be considered, and the proposed causality philosophy may be formalized and generalized for situations involving complex human environment interactions.

## Declaration of competing interest

None.

## Notes

### Competing Interest Statement

The authors have declared no competing interest.

